# Robust and time-resolved estimation of cardiac sympathetic and parasympathetic indices

**DOI:** 10.1101/2023.11.18.567211

**Authors:** Diego Candia-Rivera, Fabrizio de Vico Fallani, Mario Chavez

## Abstract

The time-resolved analysis of heart rate (HR) and heart rate variability (HRV) is crucial for the evaluation of the dynamic changes of autonomic activity under different clinical and behavioral conditions. Standard HRV analysis is performed in the frequency domain because the sympathetic activations tend to increase low-frequency HRV oscillations, while the parasympathetic ones increase high-frequency HRV oscillations. However, a strict separation of HRV in frequency bands may cause biased estimations, especially in the low frequency range. To overcome this limitation, we propose a robust estimator that combines HR and HRV dynamics, based on the correlation of the Poincaré plot descriptors of interbeat intervals from the electrocardiogram. To validate our method, we used electrocardiograms gathered from open databases where standardized paradigms were applied to elicit changes in autonomic activity. Our proposal outperforms the standard spectral approach for the estimation of low- and high-frequency fluctuations in HRV, and its performance is comparable to newer methods. Our method constitutes a valuable, robust, time-resolved, and cost-effective tool for a better understanding of autonomic activity through HR and HRV in healthy state and potentially for pathological conditions.

## Introduction

The analysis of autonomic dynamics through heart rate variability (HRV) is a standard approach for clinical and fundamental research [1–3]. Biomarkers based on HRV serve for the non-invasive analysis of physiological responses to different stimuli, which allows the assessment of several pathological conditions [4–6]. Additionally, HRV analysis can enable the characterization of neural processes, which can help to enlighten the physiological underpinnings behind homeostatic regulations, sensorimotor function, and cognition [7].

Heartbeats are generated from the continuous interactions within the autonomic nervous system, between sympathetic and parasympathetic systems [8]. These interactions occur specifically on the sinoatrial node, which contains the pacemaker cells that contract to produce the heartbeats [9]. The fluctuations in the autonomic modulations to the sinoatrial node cause the heartbeat generation at different rhythms [10], as a function of the sympathetic and parasympathetic activations that cause changes in the release rate of noradrenergic and cholinergic neurotransmitters [11]. The estimation of the sympathetic and parasympathetic activities is usually performed through HRV spectral integration at the low- (LF: 0.04–0.15 Hz) and high-frequency (HF: 0.15–0.4 Hz), respectively [12,13]. However, the spectral components of HRV series can be biased in certain conditions [14]. Specifically, the fixed subdivisions for the frequency ranges (LF and HF) cannot successfully separate the influences of the ongoing sympathetic and parasympathetic activities, which are potentially overlapped in the LF range [1]. To overcome this issue, alternative strategies have been proposed to estimate autonomic dynamics and to disentangle the low and high-frequency HRV oscillations [15–20].

We propose a method for a robust estimation of sympathetic and parasympathetic activities. The method is grounded on the measurement of the successive changes in interbeat intervals, by analyzing the changes of the Poincaré plot geometry over time. Our approach effectively estimates cardiac sympathetic and parasympathetic responses in healthy subjects. We achieve this under standard autonomic stimulation protocols, including the transition to sympathetic dominance, as achieved with postural changes and a cold-pressor test [21,22]. Additionally, we compare our estimations sympathetic activity with spontaneous fluctuations in blood pressure, a marker of sympathetic activity [23], demonstrating how effectively our method captures fluctuations in autonomic activity over time. Our method holds potential for the future development of biomarkers for clinical conditions related to dysautonomia.

## Materials and Methods

### Databases description

Twenty-four adults were recruited for the cold-pressor test. A total of 18 subjects (12 males and 6 females, aged 21 ± 1.11 years on average, body mass index 21.6 ± 1.48 Kg/m^2^ on average) were included in this study (six of them were discarded because of missing data in their ECG). Participants reported maintaining a healthy lifestyle, including non-smoking, and had no history of cardiovascular disease. Three trials of the cold pressor test were performed. Each trial consisted in a 5-minute resting period, followed by a 3-minute immersion of the hand in ice water (0°C) and a 2-minute recovery through immersion of the same hand in warmer water (32°C). Trials were considered in the -120 to 120 s with respect the onset of the cold-pressor test. Therefore, the first 3 minutes in resting state and the 2 minutes of recovery were excluded from the analysis. This approach helps minimize the potential effects on autonomic activity that might accumulate after completing one or more trials. ECG and blood pressure (reconstructed brachial arterial pressure) were measured using Finapres NOVA system (Finapres Medical Systems, Amsterdam, The Netherlands) with a sampling frequency of 200 Hz. Blood pressure was available in only 16 out of the 18 participants. For further detail on the experimental procedures, please refer to the original study [24].

Ten adults (5 males and 5 females, aged 28.7 ± 1.2 years on average, body mass index 23.7 ± 1.51 Kg/m^2^ on average) were recruited for the tilt-table test. Participants reported regularly engaging in light to moderate physical activity and had no history of cardiovascular disease. The subjects performed six trials, starting in a horizontal supine position and then transitioning to vertical position. The transitions to vertical position were performed in random order, encompassing two slow tilts (50 s from 0 to 70°), two fast tilts (2 s from 0 to 70°), and two self-paced stand ups. Subjects remain in each condition (completely horizontal or vertical position) for approximately 180 s. In this study, slow, fast and self-tilt were analyzed separately, and the two trials performed by each participant were considered as individual measurements. Trials were considered in the -120 to 120 s with respect the onset of the postural change. Therefore, the first 60 s in the horizontal position and the last 60 s in the vertical position were excluded from the analysis. This approach helps minimize the potential effects on autonomic activity that might accumulate after completing one or more trials. For further detail on the experimental procedures, please refer to the original study [25,26]. The entire protocol lasted between 55 and 75 minutes for each participant. ECG and arterial blood pressure were measured using Hewlett-Packard 1500A system (Hewlett-Packard, Palo Alto, California, United States of America) with a sampling frequency of 250 Hz.

### Estimation of sympathetic and parasympathetic indices

The R-peaks from ECGs were detected automatically using a method based on the Pan– Tompkins algorithm [27]. Consecutively, the detected R-peaks were manually corrected for misdetections. Potential misdetections were first identified by detecting peaks on the derivative of the interbeat interval series (IBI), which was computed recursively after performing manual corrections. Note that manual corrections were performed in cases of ECG borderline traces, artifacts caused by movements or contact noise, or relatively similar R and T peak amplitudes. Visual inspections of the ECG with the detected R-peaks, together with IBI histograms were consistently performed per each recording.

IBI series were constructed, based on the R-to-R-peak durations. Poincaré plot was used to depict the fluctuations on the duration of consecutive IBI [28], as shown in Figure 1A. Poincaré plots were used to depict successive differences in IBI (by plotting IBI_i_ vs IBI_i+1_, where *i* is the index of each of the IBI identified in the ECG). Poincaré plots of IBI typically depict an ellipsoid-shaped distribution, from which we quantified three features: baseline cardiac cycle duration (CCD), measured as the distance to the origin, and the variability the minor and major ratios of the ellipsoid (SD_1_ and SD_2_, respectively) representing the short- and long-term fluctuations of HRV, respectively [29]. Figure 1B displays the calculations of the Cardiac Sympathetic Index (CSI) and Cardiac Parasympathetic Index (CPI) for a single subject undergoing a cold-pressor test. These indices are derived by integrating the time-resolved estimates of CCD, SD_1_, and SD_2_. Additionally, these estimates are displayed alongside their spectral counterparts, LF and HF.

**Fig. 1.**
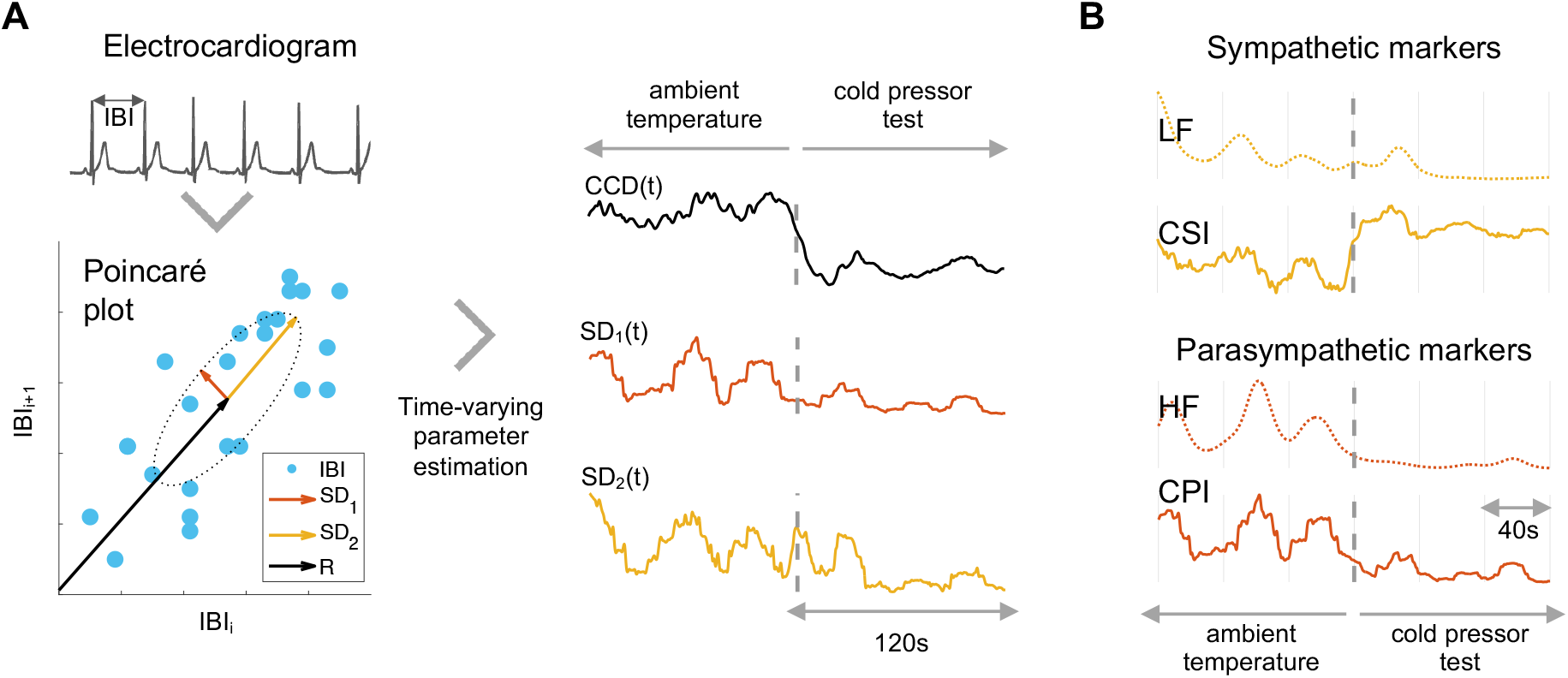
Estimation of the fluctuating parameters of the Poincaré plot. (A) The Poincaré plot illustrates the sequential changes in interbeat intervals (IBI). It allows us to estimate the Cardiac sympathetic (CSI) and parasympathetic indices (CPI) by calculating the minor (SD1) and major ratios (SD2) of the formed ellipse and the distance (R) from its center to the origin. (B) The estimated CSI and CPI are presented alongside their corresponding spectral counterparts: the low-frequency (LF) and high-frequency (HF) components of heart rate variability, which are aimed to index sympathetic and parasympathetic activity, respectively.

The time-varying fluctuations of the distance to the origin and the ellipsoid ratios were computed with a sliding-time window, as shown in Eq. 1, 2 and 3:

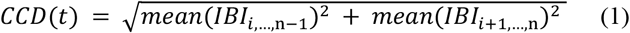

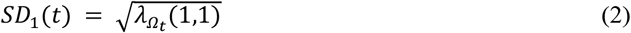

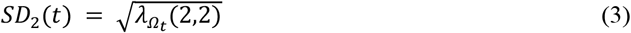

Where 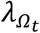 is the matrix with the eigenvalues of the covariance matrix of *IBI*_*i*,…,n−1_ and *IBI*_*i*+1,…,n_ with *Ω*_*t*_:*t* – *T* ≤*t*_*i*_ ≤*t*, and *n* is the length of IBI in the time window *Ω*_*t*_. The method implementation includes four approaches to compute SD_1_ and SD_2_: “exact”, “robust”, “95%” and “approximate”. The exact approach computes the standard covariance matrix giving the covariance between each pair of elements. The robust approach computes the covariance matrix using a shrinkage covariance estimator based on the Ledoit-Wolf lemma for analytic calculation of the optimal shrinkage intensity [30]. The 95% approach computes the covariance matrix using the Fast Minimum Covariance Determinant Estimate [31]. This covariance estimation method selects *h* observations out of total *n*. The selection of *h* fulfills the relationship *h*≈ (1 − *Outlier*) · *n*, with *Outlier* = 0.05. Then the selected points fulfill a standard covariance matrix with the lowest determinant. Finally, the approximate approach computes SD_1_ and SD_2_ as follows [16]:

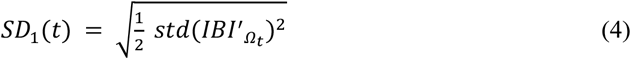

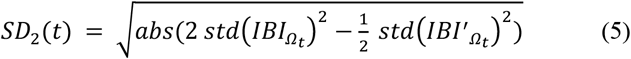

where IBI′ is the derivative of IBI, std() refers to the standard deviation, abs() refers to the absolute value, and Ω_t_: t – T ≤ t_i_ ≤ t. In this study, the main results are reported using T = 15 s, as per previous simulation studies in humans [32], and using the robust approach.

Additionally, results for the first trial of the cold pressor test are displayed for T = 5, 10, 15, 20, 25 s.

The distance to the origin *CCD*_0_ and ellipse ratios *SD*_01_ and *SD*_02._ correspond to the computation on the whole recording and were computed to re-center the time-resolved estimations of CCD, SD_1_ and SD_2_. Then, the Cardiac Parasympathetic Index (*CPI*) and the Cardiac Sympathetic Index (*CSI*), are computed as follows:

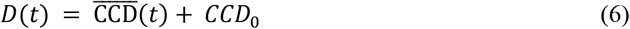

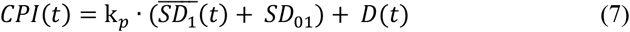

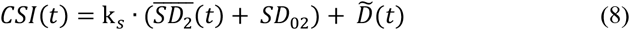

where 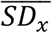 is the demeaned *SD*_*x*_ and 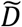 is the flipped *D* with respect the mean. The coefficients k_*p*_ and k_*s*_ define the weight of the fast and slow HRV oscillations, with respect the changes in the baseline cardiac cycle duration. The coefficients k_*p*_ and k_*s*_ define the weight of the fast and slow HRV oscillations, with respect the changes in the baseline cardiac cycle duration. Those coefficients are aimed to represent the well-stablished effects of autonomic modulations on cardiac dynamics: Sympathetic modulations primarily reduce the cardiac cycle duration [33], but also increase slower HRV changes, although, not exclusively [1,12]. Parasympathetic modulations are typically captured by quantifying overall HRV changes and the increase in cardiac cycle duration, but more specifically the faster—short-term HRV changes [1]. In this study, the values were defined empirically using one third of the datasets. For this, paired comparisons between experimental conditions were performed using a two-sided Wilcoxon signed-rank test, for k_*p*_ and k_*s*_ ranging from 1 to 10. Optimal k_*p*_ and k_*s*_ values were selected based on a good separability of the experimental conditions (assessed by the Wilcoxon’s Z), but also considering the physiological priors: dominance of the HR component in sympathetic activity and the HRV component in parasympathetic activity.

### Time-frequency estimation of cardiac sympathetic and parasympathetic activities

The estimation of sympathetic and parasympathetic activities was compared to standard HRV spectral analysis. HRV analysis in the frequency domain was computed following the adapted Wigner–Ville method for estimating the LF and HF as time series [34]. The HRV series were constructed as IBI time course. Consecutively, the IBI series were evenly re-sampled to 4 Hz using the spline interpolation [35]. In brief, the pseudo-Wigner–Ville algorithm consists of a two-dimensional Fourier transform with an ambiguity function kernel to perform two-dimensional filtering, which comprises ellipses whose eccentricities depend on the parameters *ν*_*0*_ and *τ*_*0*,_ to set the filtering degrees of time and frequency domains, respectively [36]. An additional parameter *λ* is set to control the frequency filter roll-off and the kernel tails’ size [34,36]. The parameters are set as: *v*_*0*_ = 0.03, *τ*_*0*_ = 0.06 and *λ* = 0.3, as per previous simulation studies [34]. Finally, low-frequency (LF) and high-frequency (HF) series were integrated withing the 0.04–0.15 Hz and 0.15–0.4 Hz, respectively [12,13].

The same procedure was applied to gather the LF component from QT interval series (LF_QT_), which has been proved more robust, especially for sympathetic activity estimation [19,37]. The detection of QT intervals consisted in a two-step process. From the detected R-peaks, we defined search windows around each R-peak to identify the Q peak and T peak end. The Q peak was determined by locating the minimum value within the 50 ms preceding the R-peak. For the T peak end, the minimum value within the 100-500 ms following the R-peak. Then, QT interval series were computed as the difference between the T peak end and the Q peak. From this, the time-frequency analysis was performed with an adapted Wigner–Ville method as well [34].

### Laguerre functions-based estimators of cardiac sympathetic and parasympathetic activities

The estimation of sympathetic and parasympathetic activities was compared to a model based on Laguerre expansions (namely the sympathetic and parasympathetic activity indices, SAI and PAI) [15]. This approach has been validated in numerous experimental conditions and proved valuable for modeling heartbeat generation [32]. In brief, the theoretical model expands IBI series by convolving them with a set of Laguerre functions, capturing both slow and fast fluctuations in the IBI series. Laguerre functions of orders 0 and 1 represent sympathetic oscillations, while orders 2 through 8 capture parasympathetic oscillations. Autoregressive models are used to estimate time-varying Laguerre coefficients. These coefficients are modeled as a dynamic system and estimated using a Kalman filter with a time-varying observation matrix. The kernel values for sympathetic and parasympathetic activities were derived from previously reported empirical estimates [38]. For accounting for both HR and HRV changes, the estimation of sympathetic activity is finally divided by the original IBI series, whereas parasympathetic activity is multiplied by the original IBI series. For a comprehensive overview of the methodology, please refer to the original studies [15,32,38].

### Statistical analysis

To statistically evaluate the performance of the methods in discerning the experimental conditions, we used nonparametric statistics. Time series were z-score normalized per trial. The time-resolved information for all the estimated features was condensed as the average value for each experimental session, and the group-wise descriptive measures are expressed as medians and median absolute deviations (MAD).

Paired comparisons (supine vs vertical position; ambient vs cold-pressor) were performed using a two-sided Wilcoxon signed-rank test. Significance was set to α=0.05/N, based on Bonferroni correction for N comparisons (4 comparisons for sympathetic, and 3 for parasympathetic activities).

Spearman correlations were performed to determine the relationships between CSI and blood pressure measurements. Significance was set to α=0.0001 and positive correlation coefficient (ρ). Spearman p-values were derived using a Student’s t distribution approximation. For the specific case of the tilt-table dataset, correlations were computed excluding the 0-30 s interval because of the orthostatic hypotension effect caused by the posture change [26]. To determine if the majority of participants had a significant correlation between their CSI and blood pressure series, a post-hoc statistical test was performed. This test evaluated the proportion of cases with significant correlations and calculated a p-value using the binomial cumulative distribution.

To evaluate the effect of different time window lengths (5, 10, 15, 20, and 25 seconds) on CSI and CPI computations, a group-wise statistical analysis was conducted using the non-parametric Friedman test for paired samples. This analysis was performed on data from the first trial of the cold-pressor test.

Finally, to assess sensitivity to outliers (e.g., ectopic beats), we computed the CSI on a single recording from the cold-pressor dataset. Computations were performed after artificially incorporating 30 ms delays to up to 10 IBIs. We evaluated the performance of various methods—robust, approximate, exact, and 95%—by averaging the CSI estimations from experimental conditions 1 and 2. The difference between these estimates, termed the effect magnitude, was used to quantify sensitivity. A reduced effect magnitude indicates higher sensitivity to outliers.

### Data and code availability

All physiological data used in this study are publicly available. Postural changes and intense exercise data were gathered from Physionet [25]. Cold-pressor data were gathered from Donders Institute repository [39]. Relevant code for this research work are stored in GitHub: https://github.com/diegocandiar/robust_hrv and have been archived within the Zenodo repository: https://doi.org/10.5281/zenodo.11151540 Data to reproduce the results presented in this study are available in DataDryad: https://doi.org/10.5061/dryad.6djh9w18t.

### Ethical approval statement

The cold-pressor study was approved by the local ethical committee (Radboud University, Nijmegen, The Netherlands, number ECS17022 and REC18021). The tilt-table test study was approved by the local ethical committee (Massachusetts Institute of Technology, Cambridge, United States, number COUHES 2895 and MIT CRC 512).

All participants signed informed consent to participate in the study as required by the Declaration of Helsinki.

## Results

We examined cardiac dynamics derived from HR and HRV in healthy individuals undergoing autonomic elicitation in two different conditions: tilt-table postural changes and cold-pressor test. Cardiac sympathetic and parasympathetic indices (CSI and CPI) were computed using our proposed method based on Poincaré plot descriptors of IBI series.

First, we defined the combination of HR and HRV components using the coefficients k_*s*_ and k_*p*_, which represent the weight of the HRV component in the sympathetic and parasympathetic estimations, respectively. Optimal values for k_*s*_ and k_*p*_ were determined from one-third of the datasets, based on the statistical separability of the experimental conditions. Figure 2A shows the change in the Z-value when comparing ambient temperature versus cold pressor for CSI and CPI. The results indicate that k_*s*_ = 1 provides the best separability for CSI, while CPI separability remains unaffected by changes in k_*p*_. Figure 2B shows the Z-value changes when comparing supine position versus fast tilt, exhibiting a similar trend to the cold pressor dataset. Based on these findings, we set the coefficients for the remainder of the study to k_*s*_ = 1 and k_*p*_ = 10. These values were chosen due to their alignment with well-established effects of autonomic modulation on cardiac dynamics: sympathetic modulations primarily affect baseline heart rate, while parasympathetic modulations are generally captured through HRV changes [11,15]. For illustration, Figure 2C presents the computation of CSI and CPI using k_*s*_ = 1, …, 10 and k_*p*_ = 1, …, 10.

**Fig. 2.**
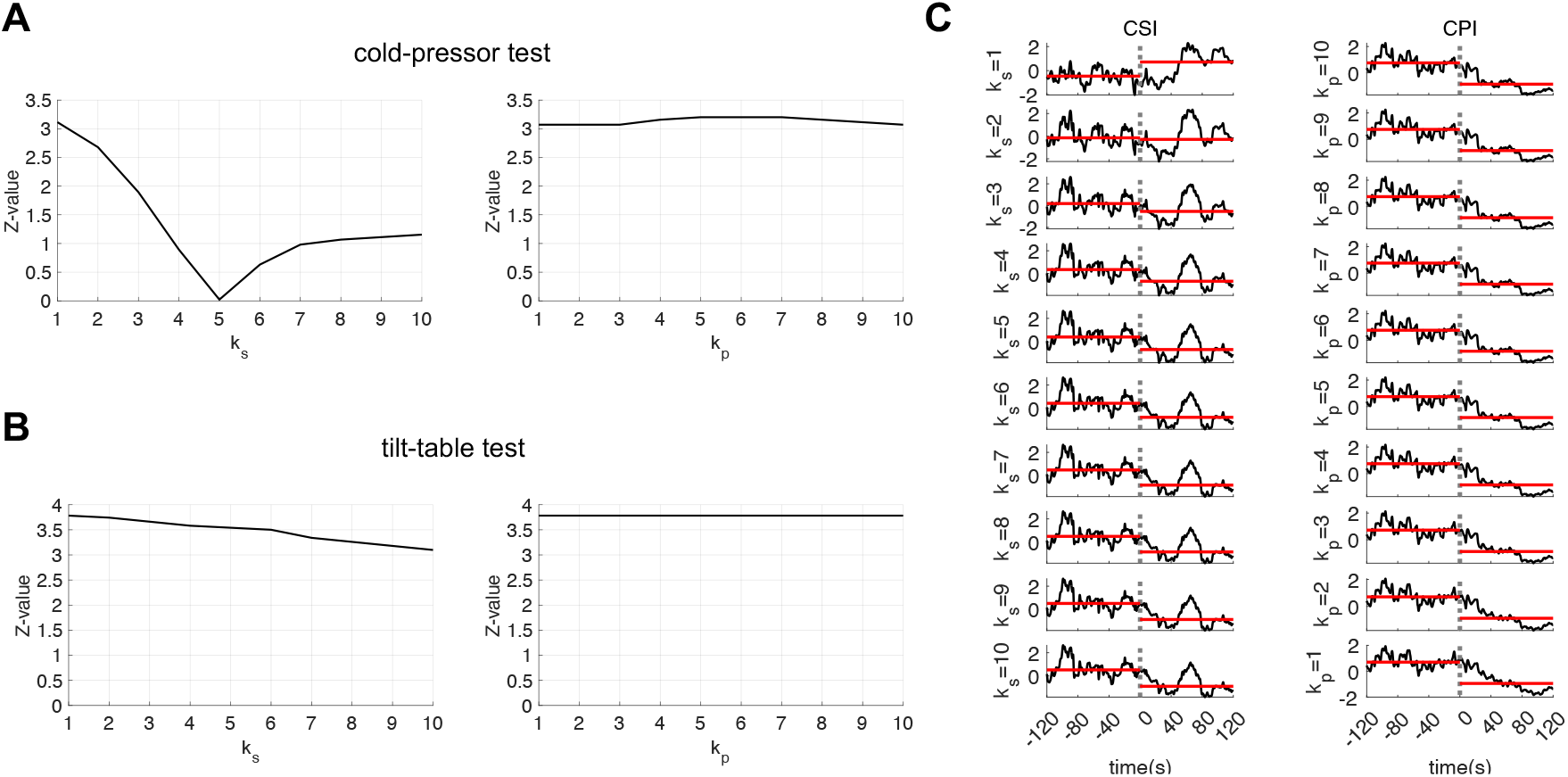
Estimation of CSI (Cardiac Sympathetic Index) and CPI (Cardiac Parasympathetic Index) for a single subject. Rows display the estimation with the HR and HRV components combined, achieved by applying weights to the HRV component using the parameters ‘k_s_’ and ‘k_p_’ for CSI and CPI estimation, respectively. The experimental condition involves the cold pressor test, with cold pressure initiating at t = 0. Horizontal red lines indicate the CSI or CPI median values before and after the cold-pressor onset

We then compared the separability of the different autonomic estimators. First, the protocol on individuals undergoing a cold-pressor test [24], in which the subjects immersed their hand in ice cold water. As shown in Figure 3A and B, our findings revealed that variations in temperature induce alterations in autonomic activity in the three trials, where an increase in sympathetic activity and a decrease in parasympathetic activity is expected [22,40–42]. While all parasympathetic markers displayed similar outcomes, the sympathetic markers exhibited divergent trends in relation to the experimental conditions. For example, LF did not show the anticipated changes in sympathetic activity, and LF_QT_ only captured the expected effects in trial 3. In contrast, the other markers, SAI and our developed CSI, performed comparably. However, while SAI estimation followed the expected trend related to changes in baseline heart rate, the HRV changes exhibited a significant high-frequency variability on top of this trend.

**Fig. 3.**
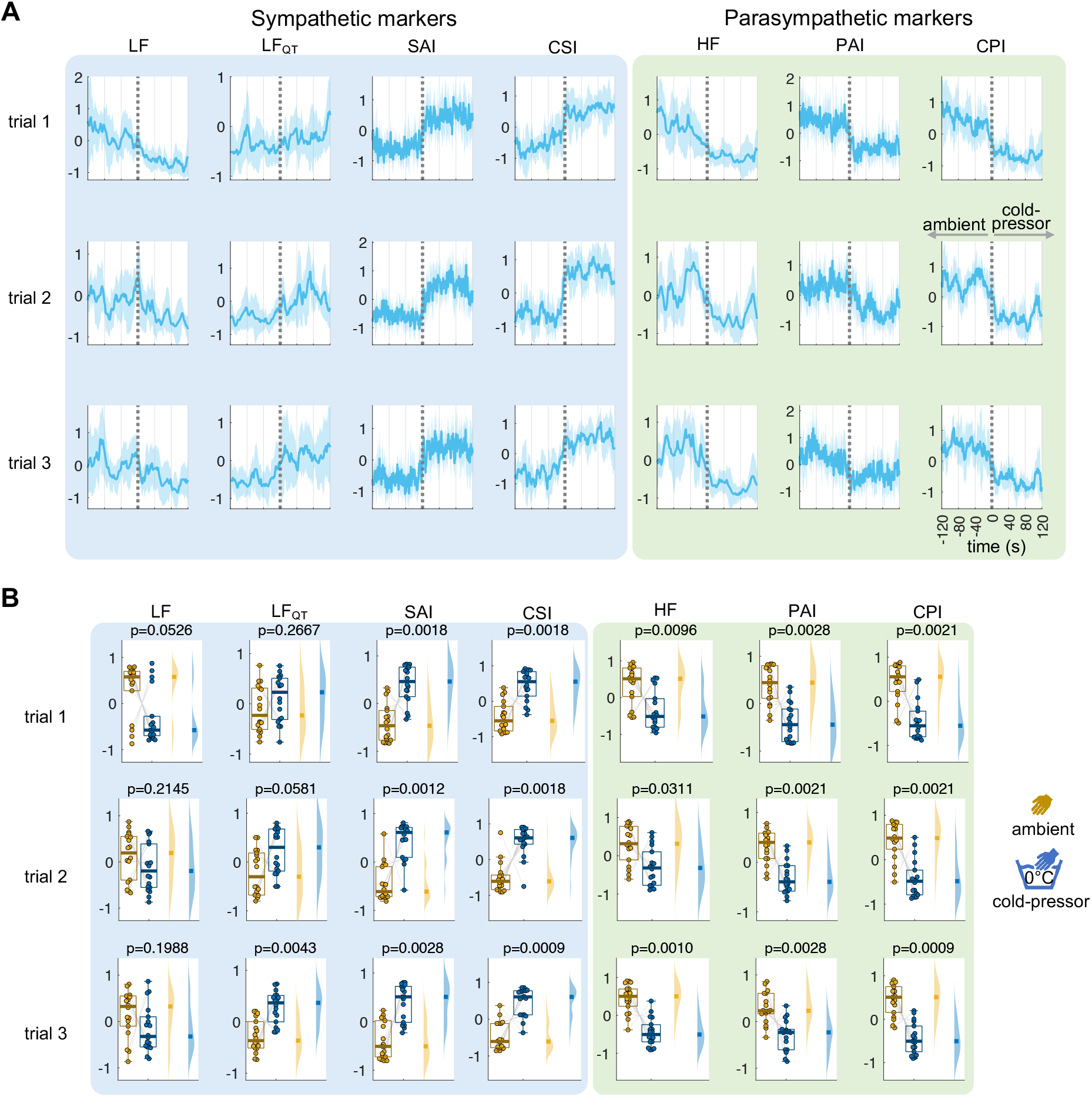
Cardiac autonomic indices CSI and CPI, and their spectral (LF, LF_QT_, HF) and Laguerre (SAI, PAI) counterparts the Cold-pressor test. The indices were used to quantify the autonomic changes triggered by the temperature changes with respect to the baseline. (A) Time course of the computed indices between -120 to 120 s with respect to the condition change onset. The plot indicates the group median and the shaded areas the median absolute deviation. Time series were z-score normalized per subject before averaging. (B) Distributions and statistical comparison using a signed rank Wilcoxon test, comparing the mean 120 s after the condition change with respect to the 120 s before. All signal in panel (A), obtained from the z-score normalization, are measured in standard deviation units.

The protocol on individuals undergoing postural changes consisted in transitioning from a horizontal/supine position to a vertical/head-up position using a tilt-table [26]. Our findings indicated that the proposed method precisely captured the dynamic fluctuations in autonomic activity in response to postural changes. Consistent with previous literature, we successfully observed the rise in sympathetic activity during the transition to an upright position [21,43–46], as depicted in Figure 4A and B. Again, when distinguishing between the two experimental conditions during postural changes, spectral counterparts underperformed, while CSI and CPI exhibited comparable performance to SAI and PAI. Similarly to the cold-pressor results, SAI and PAI present elevated high frequency components over their respective trends. Overall, these results demonstrate that the CSI and CPI estimators outperform their counterparts in these standard experimental conditions of autonomic elicitation.

**Fig. 4.**
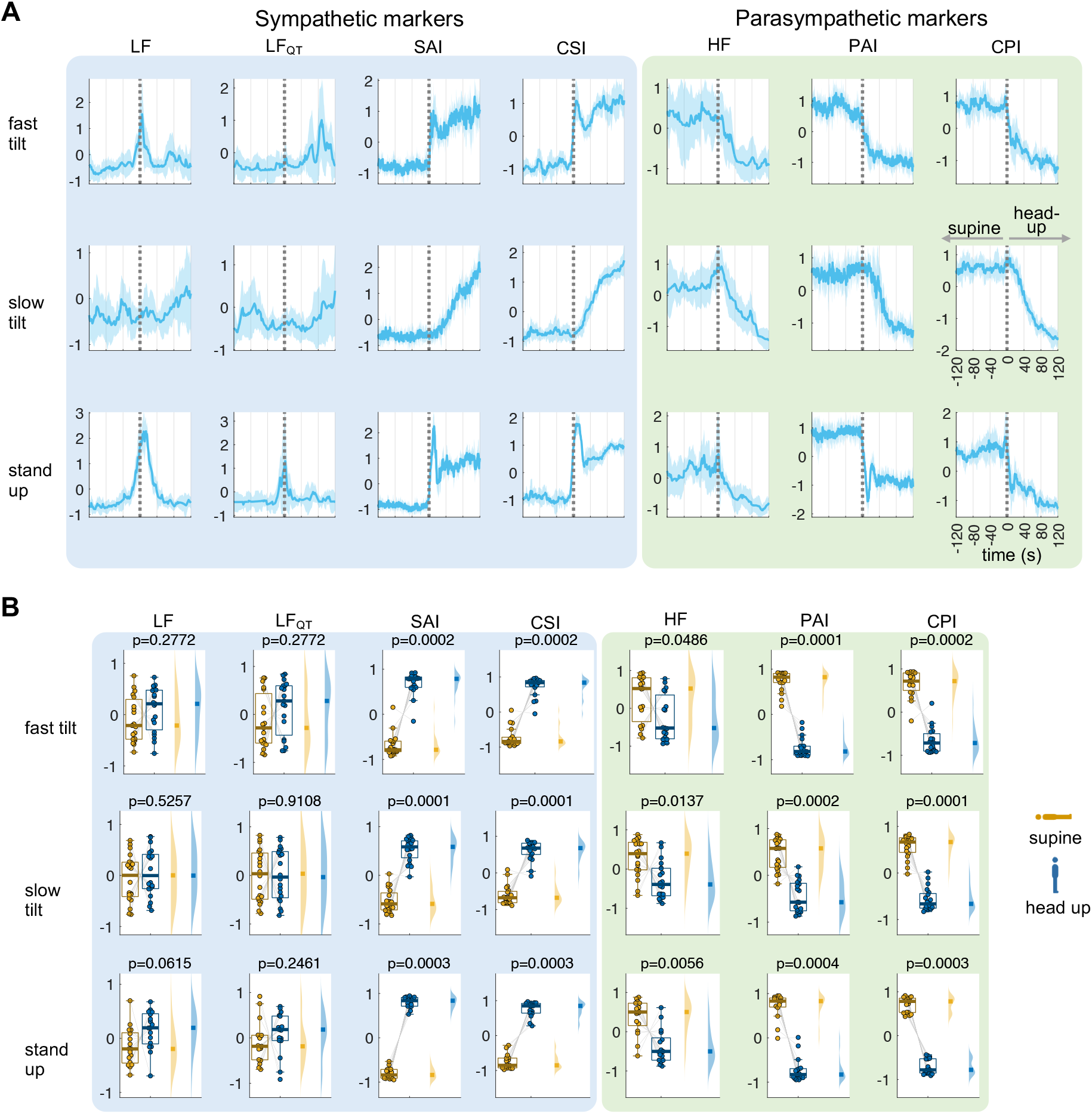
Cardiac autonomic indices CSI and CPI, and their spectral (LF, LF_QT_, HF) and Laguerre (SAI, PAI) counterparts the tilt-table test. The indices were used to quantify the autonomic changes triggered by the postural changes with respect to the baseline. (A) Time course of the computed indices between -120 to 120 s with respect to the condition change onset. The plot indicates the group median and the shaded areas the median absolute deviation. Time series were z-score normalized per subject before averaging. (B) Distributions and statistical comparison using a signed rank Wilcoxon test, comparing the mean 120 s after the condition change with respect to the 120 s before. All signal in panel (A), obtained from the z-score normalization, are measured in standard deviation units.

As an additional validation of CSI, we compared its fluctuations with concurrent blood pressure measurements. Figure 5 displays the results, showing that the cold-pressor test led to a consistent increase in both CSI and blood pressure. In the tilt table dataset, the increase in blood pressure was more subtle, with a brief reduction immediately following the position change due to the known orthostatic hypotension effect. To confirm the time-varying co-fluctuations between CSI and blood pressure, we conducted Spearman correlation analyses. Figure 5 also shows histograms, separated by experimental condition and trial, which illustrate the distribution of correlation coefficients. The majority of cases revealed a significant correlation in both the cold-pressor and tilt table datasets (all binomial tests comparing significant vs. non-significant cases, p < 0.001).

**Fig. 5.**
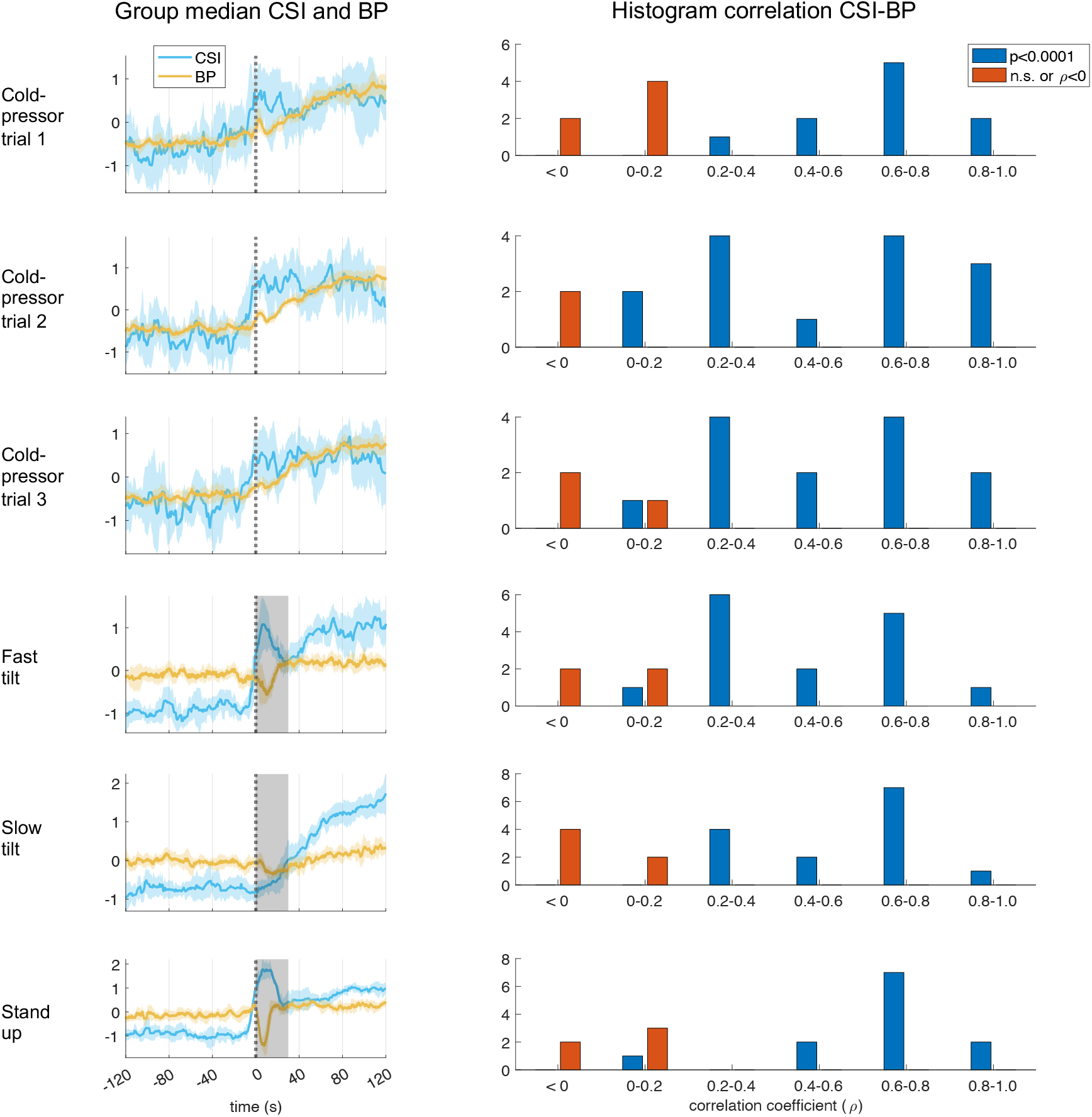
Cardiac sympathetic index (CSI) and the relationship with blood pressure (BP). The first column shows the time course of the computed CSI (in light blue), and BP (in yellow), between -120 to 120 s with respect to the condition change onset. The plot indicates the group median and the shaded areas the median absolute deviation. Gray shaded areas indicate the first 30 seconds of the tilt-table trials that were not considered due to orthostatic hypotension effects. Time series were z-score normalized per subject before averaging. All signal amplitude units are arbitrary units. The second column shows the histograms of Spearman correlation coefficients computed between CSI and BP. Significant cases (in blue) were distinguished from the non-significant ones (in red) after resulting in a positive correlation coefficient (*ρ*) and p<0.0001.

For illustration, Figure 6A presents the calculation of CSI in a subject undergoing the cold-pressor test. In this figure we highlight the impact of different time window lengths on these computations: specifically, 5, 10, 15, 20, and 25 seconds. Our findings indicate that a 5-second time window is inadequate. While it captures the well-known surge in sympathetic activity induced by cold pressure, the effect is less pronounced in the time-varying estimation compared to longer time windows. A 15-second time window or longer provides a better balance, offering both sufficient time resolution and the ability to capture gradual fluctuations in HRV around 0.1 Hz. We then statistically compared CSI and CPI using data from the first trial of the cold-pressor test. We found that the length of the time window did not significantly impact the ability to distinguish between the two experimental conditions, when comparing the statistical separability of the condition per each time window, using a paired Wilcoxon test (Figure 6B). However, a Friedman test on the CSI and CPI difference between the two conditions (cold pressor minus ambient), across different time window computations, resulted in significant differences (CSI: p= 0.0024, Friedman stat= 16.5333; CPI: p=0.000002, Friedman stat= 31.5111). These results highlight how the choice of time window can affect the assessment of modulations because the smoothing effect produced in HRV, particularly in CPI computation. It’s important to note that HRV fluctuations in the LF band can occur at frequencies as high as 0.15 Hz. Therefore, CSI estimates using a time window shorter than 7 seconds might capture HR effects rather than HRV effects. Conversely, longer time windows might reduce the visibility of HRV effects in CPI computation.

**Fig. 6.**
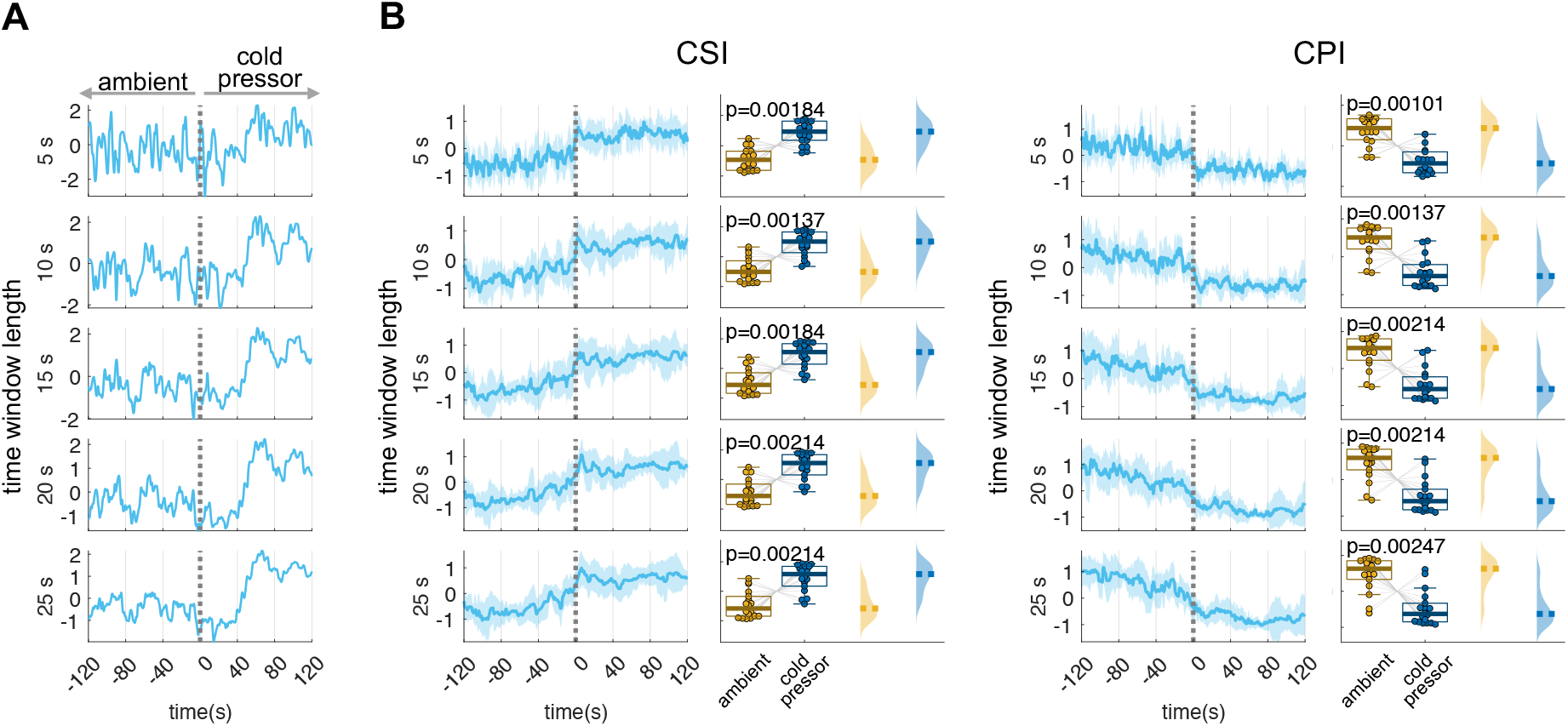
Time window length effect on the estimation of CSI and CPI. (A) Estimation of CSI (Cardiac Sympathetic Index) for one subject and (B) for the group average, for different sliding time windows. The window length displayed correspond to 5, 10, 15, 20 and 25s. The experimental condition involves the cold pressor test, with cold pressure initiating at t = 0. Statistical comparisons were performed with a Wilcoxon test between 120 s prior and after the condition change. All signals, obtained from a z-score normalization, are measured in standard deviation units.

Finally, we illustrate differences in the method implementation, which includes four approaches of computation: “exact”, “approximate”, “robust”, and “95%”. The exact approach based on the standard covariance matrix, the approximate approach based on the short-term standard deviation computations, the robust approach based on a shrinkage covariance estimator, and the 95% approach based on a 5% outliers’ rejection. The CSI estimation of one subject undergoing cold-pressure is presented in Figure 7. The estimations shown in Figure 7A correspond to the four different approaches, with each column representing the estimation results when ectopic heartbeats/outliers are externally introduced. Our findings indicate that the robust, exact, and approximate methods yield qualitatively similar estimations, with minor variations in response to the presence of ectopic heartbeats, as shown in Figure 7B. On the other hand, the 95% approach exhibits a strong resistance to outliers but results in a relatively poor overall estimation, which is demonstrated in the overall low effect magnitude (differences on CSI during the cold-pressor and baseline), but also in the high variability of the measurements with respect to the presence of outliers, as shown in Figure 7C.

**Fig. 7.**
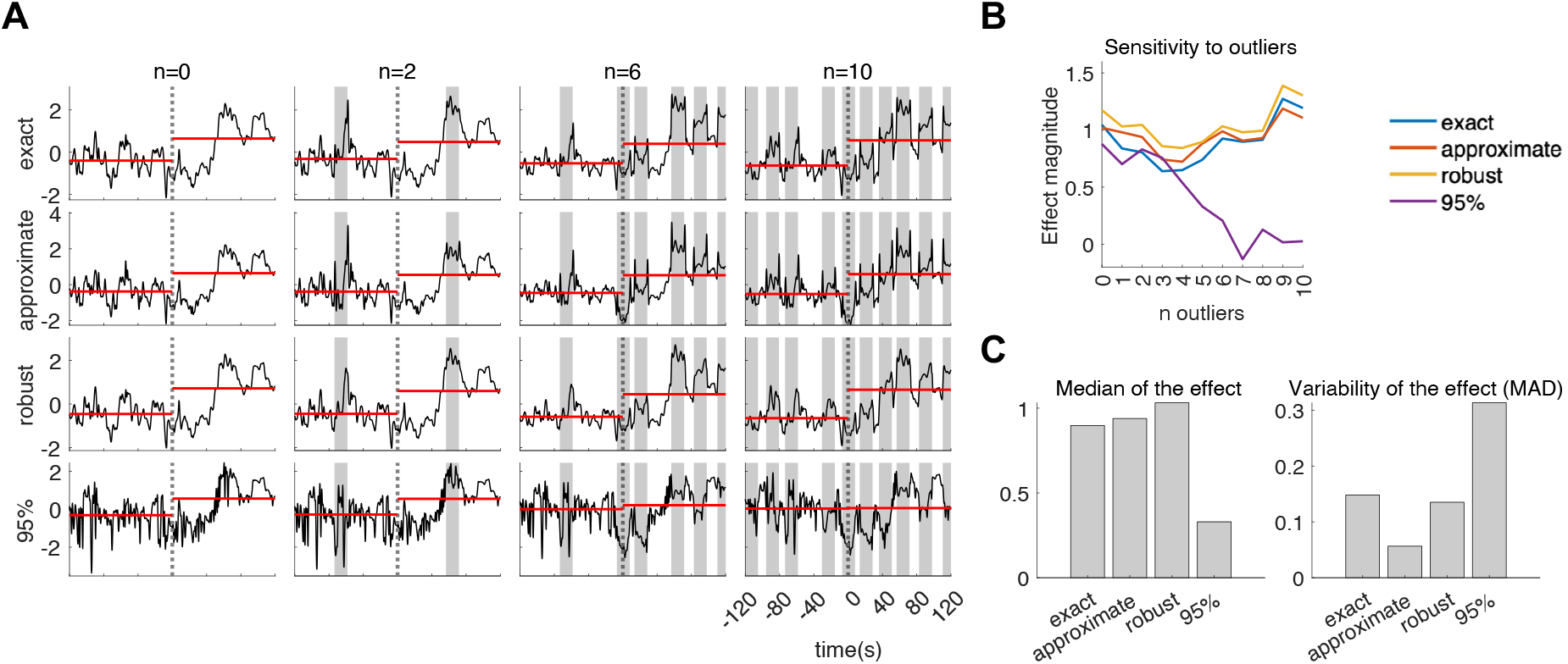
Estimation of CSI (Cardiac Sympathetic Index) for one subject. (A) Each row displays the method of estimation used (exact, approximate, robust or 95%). Each column displays the result of the estimation with the presence of misdetection of R peaks from the ECG. The number of misdetections is indicated by n = 0, 2, 6, 10. The dashed line indicates the onset of the cold-pressor at t=0. Horizontal red lines indicate the CSI median before and after the cold-pressor onset. Shaded gray areas indicate the timing in which the outliers were introduced. The misdetections were introduced by adding a delay of +30 ms to the occurrence of randomly selected R peaks. (B) Effect magnitude measured as the mean CSI during the cold pressor minus the mean CSI during ambient temperature, as a function of the number of ectopic heartbeats/outliers introduced. (C). Median and Median absolute deviation (MAD) of the effect magnitude among the eleven estimations for the number of outliers n=0,1, …, 10, for each of the four approaches.

## Discussion

We have introduced a method for the precise and time-resolved estimation of sympathetic and parasympathetic indices in humans using ECG data. Our findings highlight the remarkable consistency of the time-resolved estimations of sympathetic and parasympathetic activities across the conditions studied. This indicates that our proposed method can accurately capture and reproduce the alterations observed in sympathetic and parasympathetic activities. We also demonstrated a consistent correlation between CSI and concurrent blood pressure measurements, which further supports the validity of our method.

Our method uses Poincaré plots, which effectively depict the beat-to-beat alterations in IBIs, capturing both short-term and long-term fluctuations in HRV while also accounting for nonlinearities [28,47]. Previous studies have employed Poincaré plot-derived measures to examine sympathetic and parasympathetic influences on heart rate [16,48–50], including investigations into the changes observed in pathological conditions [51–53]. Our proposal focuses on delivering a time-resolved estimation method, enabling a comprehensive exploration of the dynamic shifts in autonomic regulations.

Our results demonstrated that CSI and CPI outperformed spectral estimations (LF, LF_QT_, and HF) and were on par with Laguerre-based estimations (SAI and PAI). One advantage of CSI and CPI is their computational efficiency, as they do not require autoregressive modeling like SAI and PAI. In the two datasets we examined, SAI and PAI showed the expected trends, but with notable high-frequency variability on top of those trends. Although we did not verify whether this variability corresponded to actual physiological dynamics, CSI and CPI provided a smoother estimator.

Physiological modeling of bodily signals plays a crucial role in uncovering the underlying aspects of autonomic dynamics by analyzing time-varying modulations of specific components. Future investigations can explore additional applications of this method, such as investigating the sympathetic and parasympathetic components involved in brain-heart dynamics [54,55], considering specific directionalities, latencies, and higher order dependencies with other physiological signals. Our proposed method, focusing on time-resolved estimations, facilitates a comprehensive exploration of dynamic shifts in autonomic regulations and their potential relation with ongoing brain activity [16,42]. This approach holds particular promise for studying pathological conditions [56] given the high level of integration in within physiological networks, which highlights the significance of modeling interoceptive processes to gain insights into multisystem dysfunctions [57]. This agrees with previous research that has already demonstrated the relevance of studying brain-heart interactions, as heartbeat dynamics have been implicated in various clinical applications [58].

Our study has some limitations, including the lack of direct ground truth measurements, such as sympathetic and parasympathetic neurograms or pharmacological manipulations [20,59]. Nevertheless, the reliability of our method is supported by tests on standardized conditions, and comparisons with blood pressure measurements—a known index of sympathetic activity—showing successful results, and therefore, serving as a validation. Another limitation of our study is the absence of a more comprehensive comparison with other measures of cardiac sympathetic and parasympathetic activity, such as symbolic representations [20]. Since the primary aim of our research is to offer a time-resolved estimation, our analysis was intentionally focused on time-resolved measures rather than a broader range of cardiac indicators. It is worth mentioning that our method relies on the geometry of the Poincaré plot, which has been criticized due to high sensitivity to the presence of outliers and artifacts [60]. To overcome this issue, we have implemented within our method a correction of potential outliers for a robust computation of the covariance matrices [30,31], which can be compared by the users to standard approaches through our open-source codes.

## Conclusion

Our method holds great potential for advancing our understanding of the dynamics of sympathetic and parasympathetic fluctuations. This tool for analyzing cardiac dynamics may also contribute to the development of physiologically inspired models for the understanding of autonomic dynamics in different contexts, such as the physiological underpinnings of sensorimotor and cognitive challenge. By employing a more accurate estimation of the ongoing autonomic dynamics we can gain deeper insights into the intricate interplay within large-scale neural dynamics.

## Acknowledgements

This research was supported by the French Agence Nationale de la Recherche (ANR-20-CE37-0012-03). DCR is supported by the European Commission, Horizon MSCA Program (grant n° 101151118).

We thank T. Heldt and colleagues for sharing the tilt-table dataset, which was supported by the NASA Cooperative Agreement NCC 9-58 with the National SpaceBiomedical Research Institute and through NlH grant Mol-RR01066 to the General Clinical Research Center at MIT.

We thank A. Mol and colleagues for sharing the cold-pressor dataset, which was supported by the Applied and Engineering Science domain (TTW) of the Netherlands Organization of Scientific Research (NWO), NeuroCIMT-Barocontrol grant 14901.

