## Supplementary Figures for "Robust and time-resolved estimation of cardiac sympathetic and parasympathetic indices"

### Supplementary Material

### Robust estimation of time-resolved sympathetic and parasympathetic outflows from heart rate variability in humans

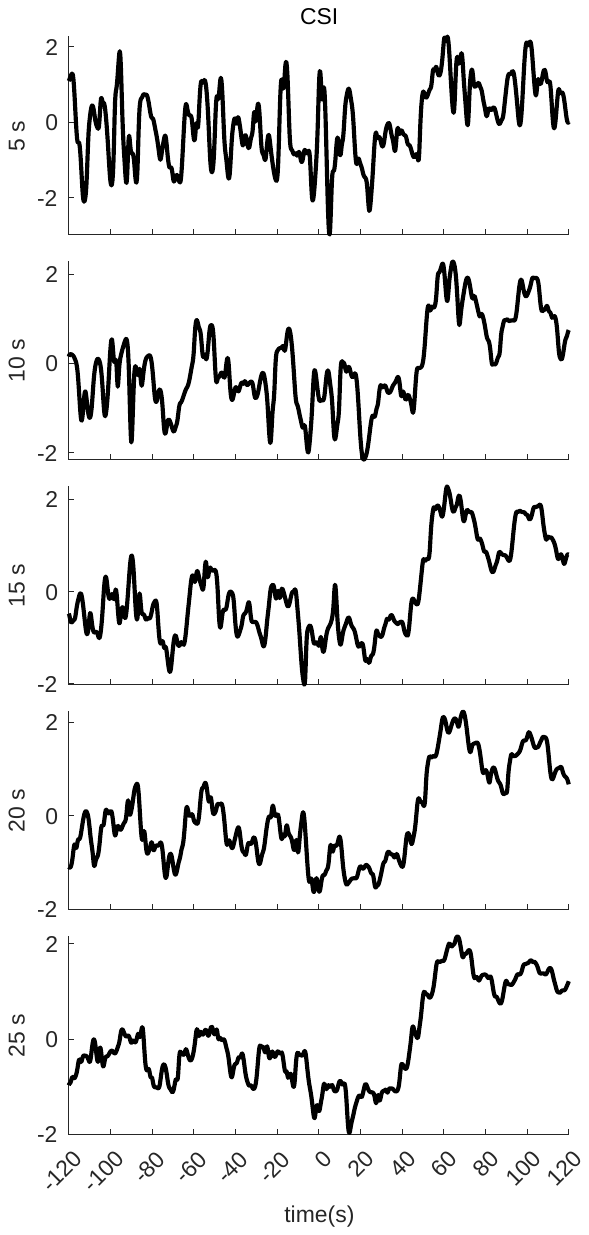

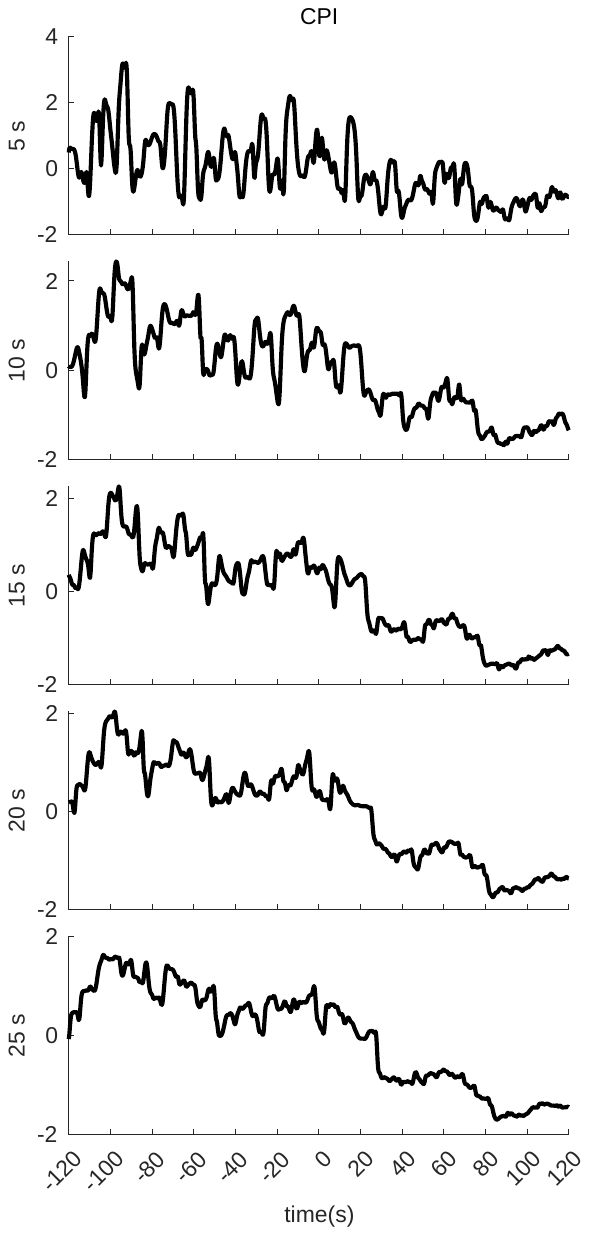

Figure 1. Estimation of CSI (Cardiac Sympathetic Index) and CPI (Cardiac Parasympathetic Index) for a single subject, for different sliding time windows. The window length displayed correspond to 5, 10, 15, 20 and 25s. The experimental condition involves the cold pressor test, with cold pressure initiating at t = 0.

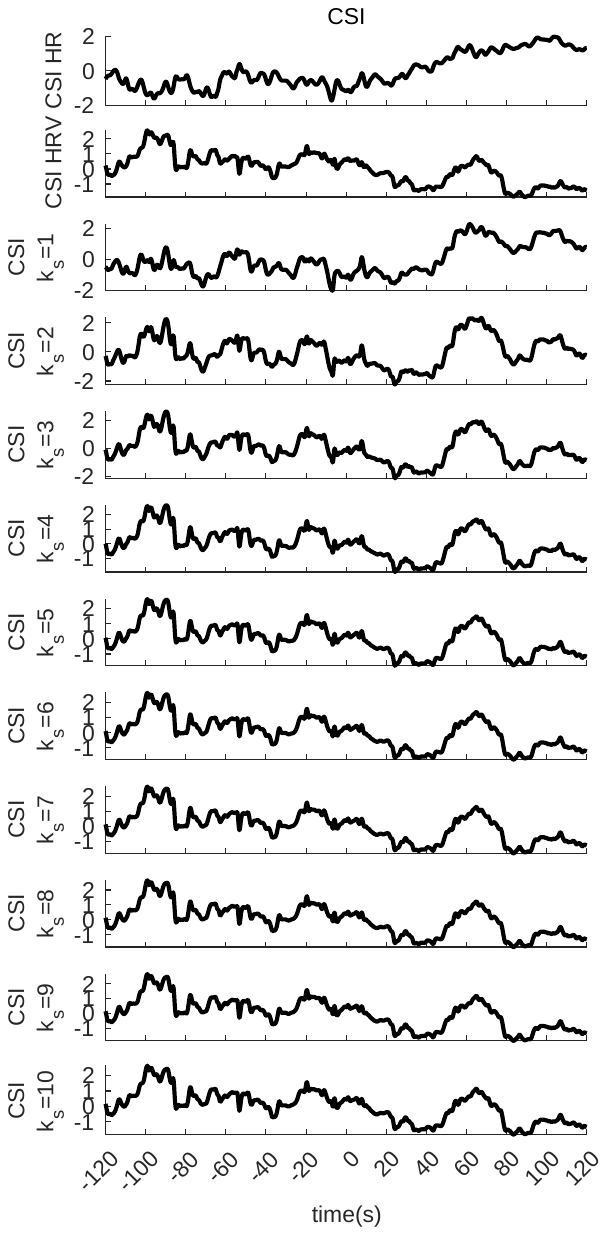

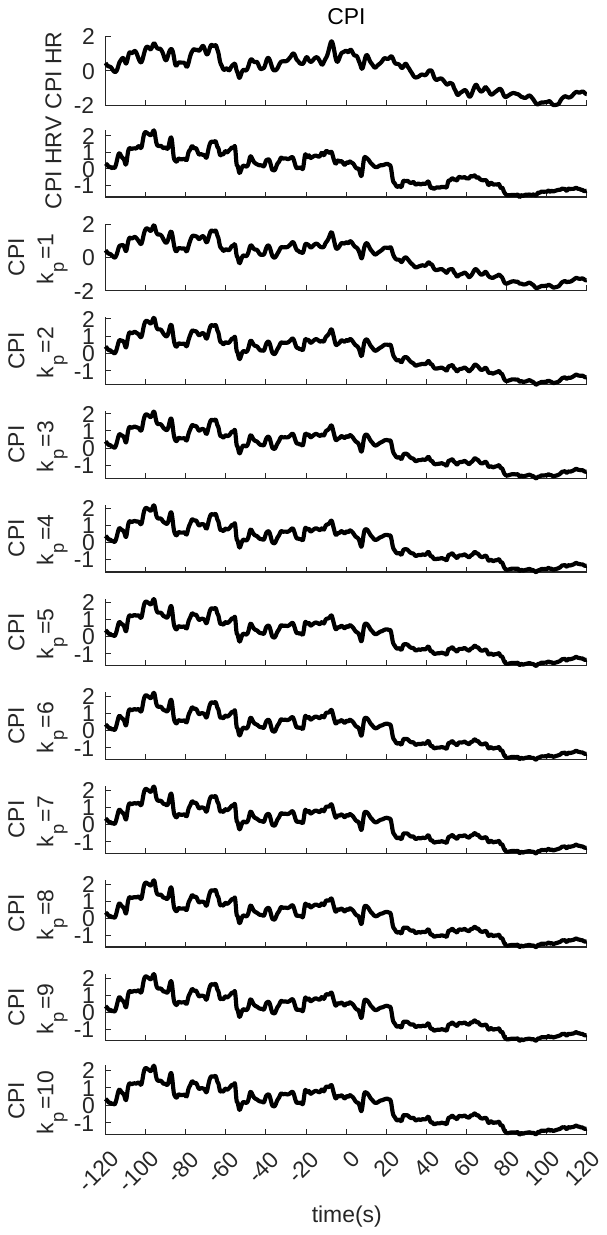

Figure 2. Estimation of CSI (Cardiac Sympathetic Index) and CPI (Cardiac Parasympathetic Index) for a single subject. The first row displays the heart rate (HR) component, while the second row the heart rate variability (HRV) component of the estimation. Subsequent rows display the estimation with the HR and HRV components combined, achieved by applying weights to the HRV component using the parameters 'ks' and 'kp' for CSI and CPI estimation, respectively. The experimental condition involves the cold pressor test, with cold pressure initiating at t = 0.

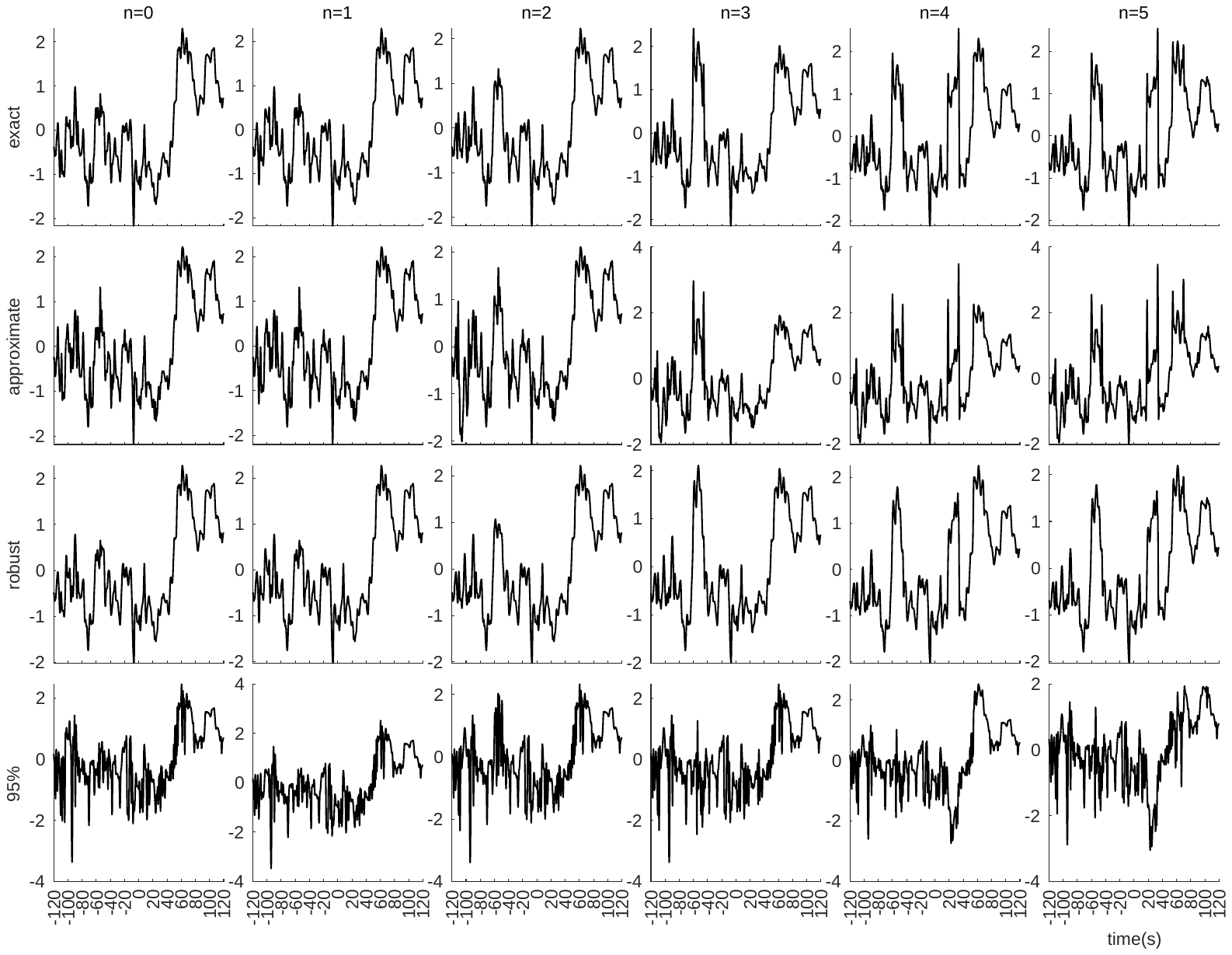

Figure 3. Estimation of CSI (Cardiac Sympathetic Index) and CPI (Cardiac Parasympathetic Index) for a single subject. Each row displays the method of estimation used (exact, approximate, robust or 95%). Each column displays the result of the estimation with the presence of misdetection of R peaks from the ECG. The number of misdetection is indicated by n = 0,…5. The misdetections were introduced by adding a delay of ±30 ms to the occurrence of randomly selected R peak. The experimental condition involves the cold pressor test, with cold pressure initiating at t = 0.
